# Changes to the resistome of *Pseudomonas aeruginosa* clone ST308 associated with corneal infection over time

**DOI:** 10.1101/2020.04.10.036673

**Authors:** Mahjabeen Khan, Mark D P Willcox, Scott A Rice, Savitri Sharma, Fiona Stapleton

## Abstract

**Objectives:** This study compared the resistomes of isolates of *Pseudomonas aeruginosa* clone ST308 from 2018 and 1997 from India.

**Methods:** Two ocular clonal type ST308 isolates of *Pseudomonas aeruginosa* (198 and 219) isolated in 2018 and five historical isolates (31, 32, 33, 35 and 37) isolated in 1997 at the LV Prasad Eye Institute in India were analysed for their susceptibilities to ciprofloxacin, levofloxacin, gentamicin, tobramycin, piperacillin, imipenem, ceftazidime and polymyxin B. DNA was extracted using the DNeasy® Blood and Tissue. Paired-end library was prepared using Nextera XT DNA library preparation kit. Libraries were sequenced on Illumina® MiSeq bench top sequencer generating 300 bp paired-end reads. Spades v3.12.0 was used for assembly, Resfinder v3.1. for acquired resistance genes and Snippy V2 for variants calling. Integron finder v1.5.1 was used to identify the integrons present in the genomes.

**Results:** The recent isolate 219 was resistant to all tested antibiotics except polymyxin while isolate 198 was resistant to ciprofloxacin, levofloxacin, gentamicin and tobramycin. Among historical isolates five were resistant to gentamicin, tobramycin and ciprofloxacin, four were resistant to levofloxacin while two were resistant to polymyxin. Twenty-four acquired resistance genes were present in the 2018 isolates compared to 11 in the historical isolates. All isolates contained the following genes encoding for aminoglycoside *aph(6)-Id*, *aph(3′)-lIb, aph(3″)-Ib)*, beta-lactam (*blaPAO)*, tetracycline (*tet(G))*, fosfomycin *(fosA)*, chloramphenicol (*catB7)*, sulphonamide (*sul1)*, quaternary ammonium (*qacEdelta1)* and fluoroquinolone (*crpP)* resistance. Isolate 198 possessed *aph(3′)-VI*, *rmtD2, qnrVC1, blaOXA-488, blaPME-1*, while 219 possessed *aadA1, rmtB, aac(6′)-Ib-cr, blaTEM-1B, blaVIM-2, mph(E), mph(A), msr(E)*. In the isolate 219 genes *blaTEM-1b*, *blaVIM-2*, *sul1*, *qnrvc1*, *rmtB* and *aadA1* were carried on class 1 integron. While an incomplete class 1 integron was also found in isolate 198 which was located on the genome where gene *rmtB*, *blaPME-1*, *qnrVC1* and *sul1* genes were positioned. There were no notable differences in the number of single nucleotide polymorphisms, but recent isolates carried more insertions and deletions in their genes.

**Conclusion:** *P. aeruginosa* ocular clonal isolates have changed over time, with strains acquiring genes and having more insertions and deletions in their chromosomal genes that confirm resistance to antibiotics.

**Highlights:**

- Recent clonal ocular isolates of *Pseudomonas aeruginosa* from India have acquired a number of resistance genes compared to historical clones
- Consequently, resistance to antibiotics particularly fluoroquinolones in recent clones of *P. aeruginosa* appears to have increased.
- The acquired resistance genes found in the recent *P. aeruginosa* isolates were related to mobile genetic elements.

## Introduction

*Pseudomonas aeruginosa* causes a variety of infections including lung infections in patients with cystic fibrosis, skin infections after burns and corneal infections (microbial keratitis). The increasing prevalence of multidrug resistant (MDR) *P. aeruginosa* reduces the treatment options and complicates management of these infections. Antibiotic resistance occurs mainly due to chromosomal gene mutations and possession of transferrable resistance determinants. [1] MDR isolates can be clonal, particularly those associated with hospital acquired infections. [2]

Clones of *P. aeruginosa* may vary based on the environments [3], and may cause infection outbreaks when these clones enter a new environment. For example, *P. aeruginosa* isolated from water sources can also be isolated from cystic fibrosis patients [4]. Only a few studies have identified clones of ocular isolates of *P. aeruginosa* [5, 6]. Five multi-drug resistant *P. aeruginosa* isolates from corneal infections have been reported to be clonal and of sequence type 308. [6] The isolates were collected in 1997 from microbial keratitis cases in India. The current study investigated the genomes of more recently collected MDR *P. aeruginosa* corneal isolates recovered from the same location in India to investigate whether this clonal variant had persisted and whether it had acquired or lost antibiotic resistance genes.

## Materials and methods

### *P. aeruginosa* genomic sequencing

DNA was extracted using DNeasy Blood and Tissue Ki (Qiagen, Hilden Germany) as per the manufacturer’s recommendations from two keratitis *P. aeruginosa* strains 198 and 219 isolated in India in 2018. A paired-end library was prepared using Nextera XT DNA library preparation kit (Illumina, San Diego, CA, USA). All the libraries were multiplexed on one MiSeq run. The raw reads of the sequenced genomes were analysed for their quality using FastQC version 0.117 (https://www.bioinformatics.babraham.ac.uk/projects/fastqc). Version 0.38 of the Trimmomatic [7] was used for trimming the adapters from the reads following *de-novo* assembly using Spades v3.13.0 [8]. Genomes were annotated using Prokka v1.12 [9]. Sequence types were investigated using PubMLST https://pubmlst.org/. Resistance genes were identified using online database Resfinder v3.1 (Centre for Genomic Epidemiology, DTU, Denmark) [10]. Mutations in the genes were detected using Snippy V2 [11] using PAO1 as a reference genome. Core genome and pan genomes were analysed using Harvest Suite Parsnp v1.2 and Roary v3.11.2 respectively. Integrons were located using Integron finder v1.5.1. The genes possessed by strains 198 and 219 were then compared to those from other ST308 isolates that had been previously examined [6].

### Antibiotic resistance

Strains 198 and 219 (isolated in 2018) and five strains isolated in 1997 PA31, PA32, PA33, PA35 AND PA37 [6] were screened for resistance to a variety of antibiotics which are commonly used to treat microbial keratitis [12]. The minimum inhibitory concentration (MIC) and minimum bactericidal concentration (MBC) of ciprofloxacin, levofloxacin, gentamicin, ceftazidime (Sigma-Aldrich, St. Louis Missouri, USA), polymyxin B (Sigma-Aldrich, Vandtårnsvej, Søborg, Denmark) tobramycin, piperacillin (Cayman Chemical Company, Ann Arbor, Michigan, USA) and imipenem (LKT Laboratories Inc, Minnesota, USA) were determined using the broth microdilution method in 96-wells plates following CLSI guidelines. The concentrations of antibiotics tested ranged from 5120 µg/ml to 0.25 µg/ml. The susceptibility results were interpreted using EUCAST v9 [13] and CLSI [14] breakpoints for antibiotics.

## Results

### Antibiotic susceptibility and Sequence type analysis

Isolates 198 and 219 had sequence type 308 indicating that these strains were clonally related to the ST308 strains isolated in 1997 at the same hospital.

Isolate 219 was resistant to all antibiotics (Table 1) other than polymyxin (MIC= 0.5 µg/ml, MBC=1 µg/ml). Isolate 198 was resistant to ciprofloxacin, levofloxacin, tobramycin and gentamicin, and showed intermediate susceptibility to polymyxin (Table 1). The isolates from 1997 were all resistant to gentamicin and tobramycin and showed intermediate or definite resistance to imipenem (Table 1). All five isolates from 1997 were resistant or had intermediate resistance to ciprofloxacin and four were resistant to levofloxacin (Table 1). Two isolates from 1997 showed intermediate resistance to polymyxin (Table 1). Overall, the MIC and MBC values to ciprofloxacin and levofloxacin of 198 and 219 were higher than those recorded for the historical isolates (Table 1).

**Table 1.**
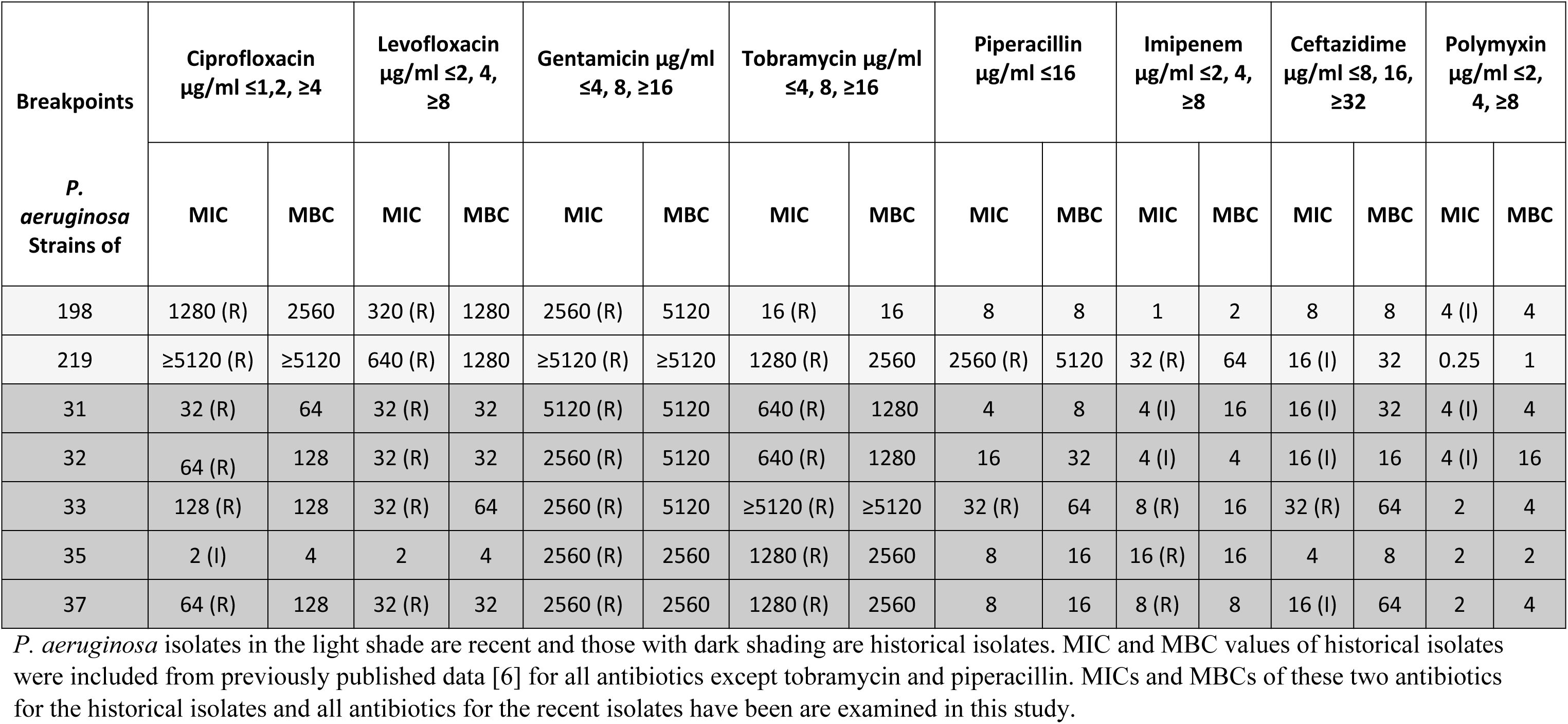
Antibiotic susceptibility of *P. aeruginosa* isolates

### Possession of horizontally-acquired resistance genes

In total 24 acquired resistance genes were present in the ST308 isolates of *P. aeruginosa* (Table 2). The isolates from 1997 all possessed the same 11 resistance genes. However, the isolates from 2018 had acquired additional resistance genes. Isolate 198 carried 15 and 219 carried 20 resistance genes. Ten resistance genes were common to all seven isolates (Table 2). These ten genes were three aminoglycoside resistance genes *(aph(6)-Id*, *aph(3′)-lIb, aph(3″)-Ib)*, a beta-lactam resistance gene (*blaPAO)*, a tetracycline resistance gene (*tet(G))*, a fosfomycin resistance gene *(fosA)*, a chloramphenicol resistance gene (*catB7)* a sulphonamide resistance gene (*sul1)*, and a quaternary ammonium compound resistance gene (*qacEdelta1)*. The recent isolates lacked one beta lactam gene (*blaOXA-50)* which was present in all the historical isolates.

**Table 2.**
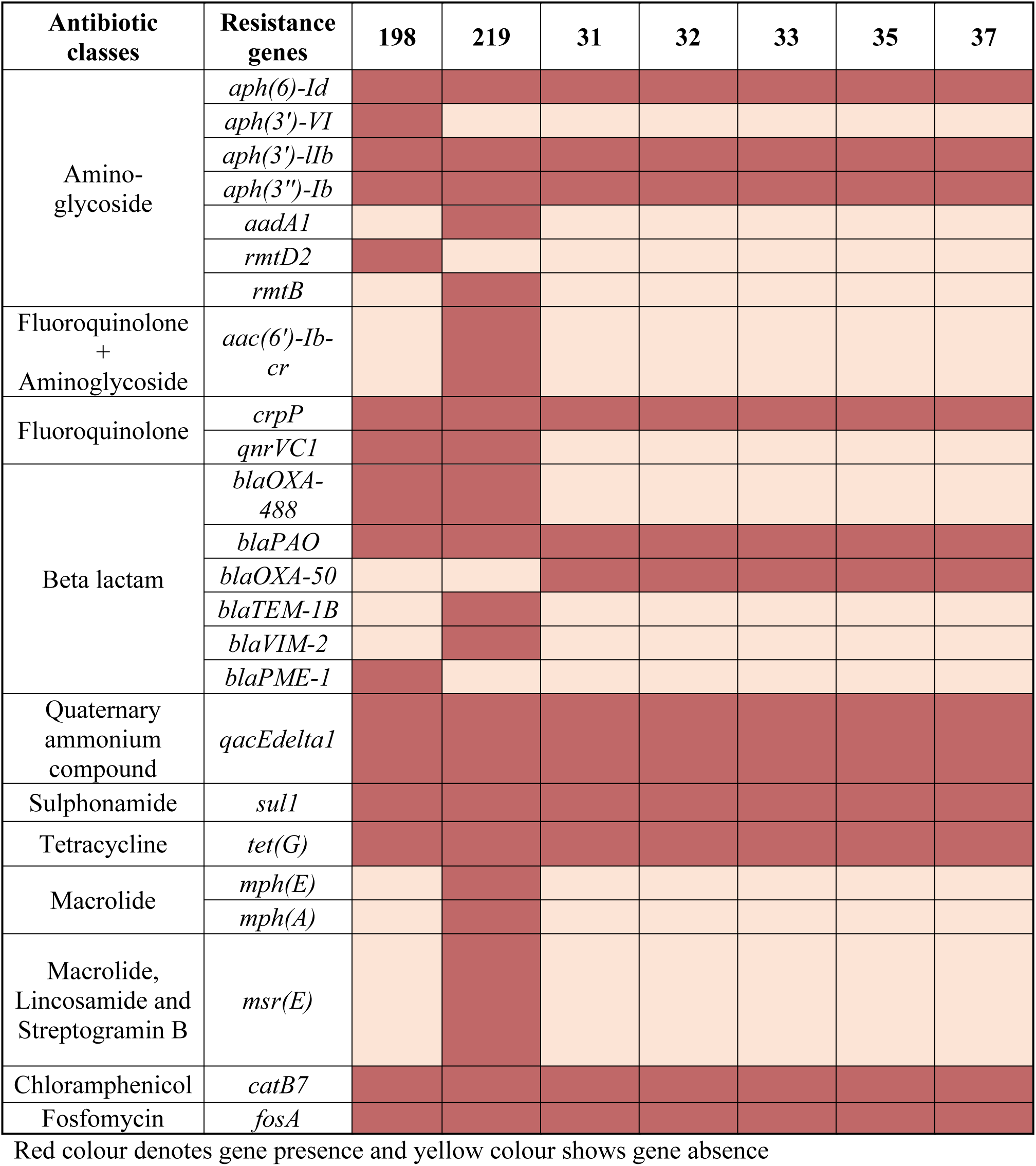
Presence of acquired antibiotic resistance genes in *P. aeruginosa* ocular isolates.

### Aminoglycoside resistance genes

Strain 198 had acquired a 16S rRNA methylase gene *(rmtD2)* carried on class 1 integron and three aminoglycoside modifying enzyme genes *aph(6)-Id*, *aph(3′)-lIb*, and *aph(3″)-Ib)*. Strain 219 had acquired three different aminoglycoside resistance genes, a 16S rRNA methylase (*rmtB)*, a streptomycin adenylyltransferase gene *(aadA1)* carried on class 1 integron and an aminoglycoside acetyltransferase gene *(aac(6′)-Ib-cr).* Strain 219 had also acquired the plasmid related aminoglycoside and fluoroquinolone resistance gene *aac(6′)-Ib-cr*.

### Fluoroquinolones resistance genes

One fluoroquinolone resistance gene, *crpP*, was present in all isolates from 1997 and 2018. An integron-related fluoroquinolone resistance gene *qnrVC1* was only present in isolates from 2018 (Table 1) and was carried on class 1 integron on both isolates 198 and 219. The plasmid related aminoglycoside and fluoroquinolone resistance gene *aac(6′)-Ib-cr* was found in strain 219 and this strain had higher MICs for ciprofloxacin and levofloxacin compared to strain 198 and the 1997 isolates.

### Beta-lactam resistance genes

The metallo-beta-lactamase gene class B metallo-b-lactamase *blaVIM-2* and a transposon (Tn2) encoded gene *blaTEM-1B* had been acquired by isolate 219 and were carried on class 1 integron. An extended spectrum plasmid-related class A beta lactamase gene *blaPME-1* had been acquired by 198 and were carried on class 1 integron. A class-D beta lactamase gene *blaOXA-488* had been acquired by both 198 and 219.

### Non-synonymous mutations in the ST308 resistome

Table 3 details the non-synonymous mutations leading to changes in the nucleic acid sequence in the resistance genes of these *P. aeruginosa* isolates, including those related to efflux pumps, antibiotic-inactivating enzymes and drug target alterations. These non-synonymous mutations were made in comparison to the reference genome of strain PAO1. The number of mutations in almost all of the genes remained same in the 1997 and 2018 isolates. However, the efflux pump gene *opmH* contained 10 SNPs in the isolates from 2018, but only 1-5 SNPS in the isolates from 1997 (Table 3). Similarly, *oprD* also contained 8 SNPs in the two 2018 isolates. Furthermore, non-synonymous insertions/deletions [15] and frame-shift mutations were found in the two isolates from 2018 (Table 4) whereas the ST308 isolates from 1997 had no insertions/deletions or frame-shift mutations [6].

**Table 3.**
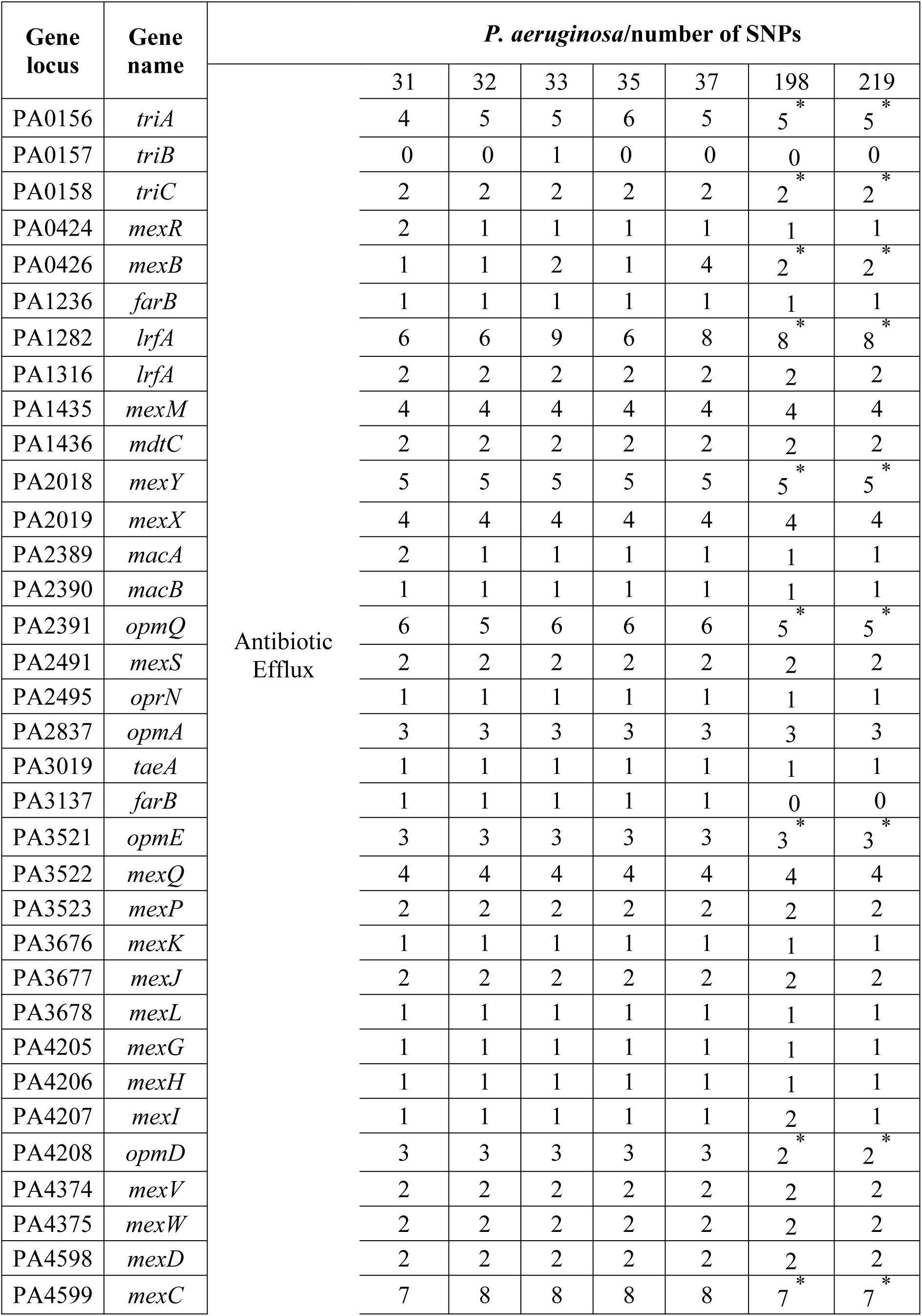
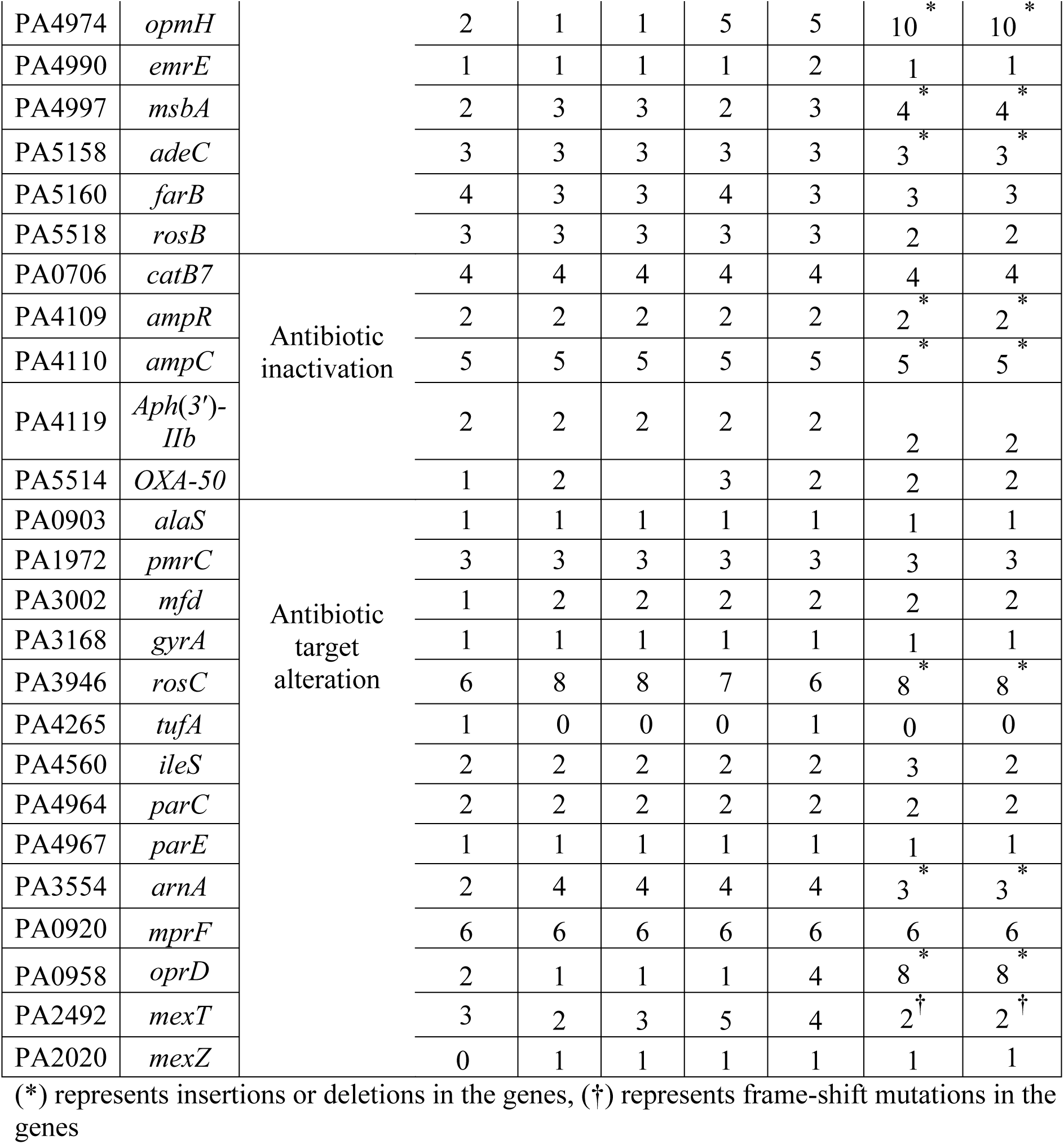
Single nucleotide polymorphism due to non-synonymous mutations in the genes of *P. aeruginosa* genes

**Table 4.** Insertion/deletion and frame-shift mutation in the resistance genes of the 2018 isolates.

| Gene locus | Genes | Number of deletions and insertions in the genes |  |
| --- | --- | --- | --- |
|  |  | 198 | 219 |
| PA0156 | *triA* | 2 | 2 |
| PA0158 | *triC* | 2 | 1 |
| PA0426 | *mexB* | 1 | 1 |
| PA1282 | *lrfA* | 2 | 2 |
| PA2018 | *mexY* | 1 | 1 |
| PA2391 | *opmQ* | 3 | 3 |
| PA3521 | *opmE* | 1 | 1 |
| PA4208 | *opmD* | 1 | 1 |
| PA4599 | *mexC* | 1 | 1 |
| PA4974 | *opmH* | 7 | 8 |
| PA4997 | *msbA* | 2 | 2 |
| PA5518 | *adeC* | 1 | 1 |
| PA4109 | *ampR* | 1 | 1 |
| PA4110 | *ampC* | 2 | 2 |
| PA3946 | *rosC* | 1 | 1 |
| PA3554 | *arnA* | 2 | 2 |
| PA0958 | *oprD* | 6 | 6 |
| PA2492 | *mexT* | 1Frame-shift deletion | 1Frame-shift deletion |

### Phylogeny of ST308 isolates

The genomes of these ST308 isolates were aligned using PAO1 as a reference for the core genome and pangenome phylogeny. In the core genome phylogenetic analysis, all the isolates were clustered together in a single group. The number of core genes and total genes were same for all isolates except isolate 219 which had a larger number of total genes but a similar number of core genes to all other isolates (Table 5).

**Table 5.** Number of genes present in Core and pan genomes of *P. aeruginosa* isolates.

| *P. aeruginosa* isolates | Core genes | Total genes |
| --- | --- | --- |
| 31 | 5445 | 6937 |
| 32 | 5447 | 6927 |
| 33 | 5440 | 6932 |
| 35 | 5442 | 6932 |
| 37 | 5450 | 6958 |
| 198 | 5454 | 6882 |
| 219 | 5451 | 7247 |

## Discussion

This study examined whether the resistome of the ST308 clone of ocular isolates of *P. aeruginosa* had changed over time. Previously, *P. aeruginosa* ST308 clones had been reported as multidrug-resistant isolates of nosocomial [16] ocular [6] and canine origin [17]. For the *P. aeruginosa* isolates from three different sources the MIC to imipenem was high which was similar to the finding in the ocular isolates of *P. aeruginosa* in the present study. The ocular isolates of clone ST308 from 2018 had acquired additional resistance genes and had changes in the mutational patterns of the resistance genes compared to ocular isolates from 1997.

Two different variants of 16S rRNA methylase, *rmtD2* and *rmtB*, related to aminoglycoside resistance were found in the 2018 isolates. These genes have not been reported previously in ST308 but other variants of the same genes have been identified in clones ST316 and ST235 [18] the latter clone being identified as a widespread multi-drug resistance clone. The presence of a larger group of beta lactam resistance genes, specifically those acquired on mobile genetic elements including class A and B metallo-beta lactam genes including *blaTEM-1B, blaVIM-2, blaPME-1* carried on integron is a unique finding related to ST308 in the current study. These beta lactam resistance genes have not been reported previously in strains of this clone [6]. However, the possession of *sul1* gene in isolates of present study was similar to the similar ST308 found previously [17]. The possession of *blaVIM-2* and *blaTEM-1B* may have been responsible for the high MIC to piperacillin and imipenem of PA219. Previously, these genes were associated to increased MIC of imipenem and piperacillin/tazobactam in *P. aeruginosa* isolates [19].

Metallo-beta lactam genes are usually found on class-1 integrons along with other antibiotic resistance determinants [20] which is similar to the present study but identification of class 1 integron carrying resistance genes in the ocular *P. aeruginosa* isolates is a novel finding. These metallo-beta lactam genes are easily transmissible on mobile genetic elements such as transposons, plasmid-integrative conjugative elements and genomic islands. These metallo-beta lactam genes (*blaTEM-1B, blaVIM-2, blaPME-1*) have not been previously reported in ST308 but have been found in ST111 and ST235 [21] [22]. Although different variants of these genes were found in the similar ST308 before [16, 17]. Acquired genes within the mobile genetic elements of ST308 clones were not been identified in an earlier report [6]. Both recent isolates 198 and 219 had acquired genes associated with mobile genetic elements in the current study.

The presence of the plasmid related fluoroquinolone resistance gene *qnrVC1* [23] and the recently reported plasmid related gene *crpP* [24] are also novel findings in the current study related to clonal ST308 *P. aeruginosa* isolates. All isolates contained the fluoroquinolone **r**esistance gene *crpP*, but this had not been identified as a potential plasmid related fluoroquinolone resistance gene prior to the publication of resistance genes of the 1997 isolates [6]. Usually fluoroquinolone resistance is due to mutation in DNA gyrase and topoisomerase IV genes [25]. However, in the 2018 isolates of ST308 very high MICs to ciprofloxacin and levofloxacin might be due to the acquisition of *qnrVC1*. Strain 219 had also acquired the plasmid related fluoroquinolone resistance gene *aac(6′)-Ib-cr* [26] which can confer resistance to both fluoroquinolones and aminoglycosides [27]. Previously this gene was found responsible for the 16 to 128-fold higher MICs for ciprofloxacin in the transconjugants bacteria of family Enterobacteriaceae [28] and MIC of 64 µg/ml of ciprofloxacin to MDR *P. aeruginosa* isolates [29]. These additional resistance imposing elements to fluoroquinolones suggest that alternative treatments for keratitis other than fluoroquinolone monotherapy should be considered. Acquisition of larger number of aminoglycoside and beta lactam resistance genes is alarming because, where first line therapy such as monotherapy with fluoroquinolones fails, fortified antibiotics [30] such as gentamicin plus cephalosporins are often prescribed.

Among all the *P. aeruginosa* isolates, the core genome was composed of almost similar number of genes which was perhaps indicative of the collinear nature of conserved genome of *P. aeruginosa* isolates. [31, 32] However, a larger pan genome of isolate 219 indicates greater genomic diversity due to acquisition of genes from the same or different species or genera. This fact might relate to the larger number of acquired genes in isolate 219 by horizontal gene transfer. [31] Identification of indels (insertion/deletion polymorphisms due to non-synonymous mutations) in the 2018 keratitis isolates of *P. aeruginosa* which were not present in the strains isolated in 1997 [6] as well as the increased presence of certain SNPs suggest that there was an increase in selection pressure in the environment that has selected for these mutations.

Increases in the resistance of keratitis isolates to the fluoroquinolone moxifloxacin have been associated with an increase in average diameter of the infiltrate or scar, a slower time to re-epithelialization and decrease in final visual acuity. [33, 34] Therefore, the findings from the current study showing that strains of *P. aeruginosa*, at least in this Indian environment, have gained additional resistance genes and higher levels of resistance suggests that treatment of keratitis might be becoming more problematic.

### Nucleotide accession

The nucleotide sequences are available in the GenBank under the Bio project accession number PRJNA590804.

## Acknowledgements

The authors would like to acknowledge the Singapore Centre for Environmental Life Sciences Engineering (SCELSE), whose research is supported by the National Research Foundation Singapore, Ministry of Education, Nanyang Technological University and National University of Singapore, under its Research Centre of Excellence Programme. Sequencing of DNA was carried out with the help of Stephen Summers using the sequencing facilities at SCELSE. We are also thankful to UNSW high performance computing facility KATANA for providing us cluster time for data analysis.

## Authors’ contributions

MW: Conceptualization of the study, manuscript review and editing.

MK: Experimental procedures, genome analysis and writing of the manuscript.

FS: Conceptualization of the study, manuscript review and editing.

SS: Donation of strains and manuscript review.

SR: Genome sequencing facilitation, manuscript review. All authors have approved the final article

